# marge: An API for Analysis of Motifs Using HOMER in R

**DOI:** 10.1101/249268

**Authors:** Robert A. Amezquita

## Abstract

Profiling of open chromatin regions through the assay for transposase-accessible chromatin (ATAC) and transcription factor binding via chromatin immunoprecipitation (ChIP) sequencing has increased our ability to resolve patterns of putative transcription factor binding sites at the genome-wide level. Popular tools such as [HOMER (http://homer.ucsd.edu/homer/) and [MEME] (http://meme-suite.org/) have driven forward the analyses of sequence composition, deriving *de novo* motifs and searching for the enrichment of known motifs. However, their interfaces do not allow for the construction of parallel inquiries of multiple datasets. Furthermore, their results do not conform to formats amenable to ‘tidy’ analyses, presenting a significant bottleneck in motif analysis. Here, I present ‘marge’, a companion ‘R’ package that interfaces with HOMER to facilitate the construction of queries and to tidy results for further downstream analyses.

## Introduction

Profiling of open chromatin regions through the assay for transposase-accessible chromatin (ATAC) and transcription factor binding via chromatin immunoprecipitation (ChIP) sequencing has increased our ability to resolve patterns of putative transcription factor binding sites at the genome-wide level. Popular tools such as HOMER and MEME have driven forward the analyses of sequence composition, deriving *de novo* motifs and searching for the enrichment of known motifs. However, their interfaces do not allow for the construction of parallel inquiries of multiple datasets. Furthermore, their results do not conform to formats amenable to ‘tidy’ analyses, presenting a significant bottleneck in motif analysis. Here, I present `marge`, a companion R package that interfaces with HOMER to facilitate the construction of queries and to tidy results for further downstream analyses.

## Methods

### Implementation

`marge` is an `R` package that makes use of various `tidyverse` conventions, allowing for modern conventions of R programming and manipulations. `marge` interfaces directly with HOMER to construct queries, execute motif enrichment searches, and organize results. While this requires a separate installation of HOMER and introduces this long-term dependency, core HOMER routines have been stable for a long time, with infrequent updates. Given the popularity of HOMER, the syntax and construction of queries will thus be familiar to the audience utilizing the `marge` companion package. However, to conform to modern standards of software design, certain core routines, such as `findMotifsGenome.pl` which encompasses many different tasks, have been partitioned into distinct R functions. For example, motif analysis and motif location identification have been split up into distinct functions `find_motifs_genome` and `find_motifs_instances`, respectively.

### Operation

`marge` is distributed on Bitbucket as an `R` package, and is compatible with Mac OS X, Windows, and Linux. The latest version of HOMER (v4.9 as of this writing) should be installed prior to `marge`, and configured with the desired genomes and feature sets. Subsequently, `marge` can be installed from the repository. R Package dependencies and system requirements are documented in the repository and automatically installed in conjunction with the marge package. Online documentation is also available at `marge.aerobatic.io`.

### Use Cases

To demonstrate the functionality and utility of `marge`, we present a tutorial of core functionality and present potential use cases for high-throughput motif enrichment studies and subsequent downstream analyses.

### Conventions

Code objects shall be referred to in-line with the code font, whereas R functions will be referred to with a pair of parentheses following the function name, as in `find_motifs_genome()`. `R` code chunks will be presented as a block. Comments describing relevant portions of code are prepended by hashes (`#`), and all else is runnable `R` code. Results of code chunks are printed in separate block below, with each line of output prepended by a double hash (`##`). An example of code chunk and output formatting follows:

~~~
salutation <- c('hello world')
salutation *# view output*

## [1] "hello world"
~~~

Lastly, documentation for functions can be accessed invoking through help utility in R. Online documentation with the same content is also available under the ‘Reference’ tab.

### Preamble

The latest version of HOMER should be installed prior to installing `marge` as documented online in the introduction/install section. Following installation, HOMER packages should additionally be installed, which contains the necessary sequence data to perform motif enrichment, such as those that follow:

~~~
## In a terminal/command-line
perl /path-to-homer/configureHomer.pl-install hg38 *# human sequence data*
perl /path-to-homer/configureHomer.pl-install mm10 *# mouse sequence data*
~~~

Additionally, users should take care to ensure that HOMER is added to the executable path, such that entering `findMotifsGenome.pl` on the command-line works as expected. For further details, we again refer to the introduction/install section of the HOMER documentation.

### Basic Usage

What follows is a basic tutorial of a typical workflow in performing motif enrichment analysis from a given, single set of regions. This section will introduce the various verbs available in `marge` for this task.

### Check HOMER and Installed Packages

To check that HOMER was installed properly and is detected by `marge` invoke the `check_homer()` function. Sequence annotation data for the genome of interest should also have been installed prior to running HOMER or `marge`. To check what is currently available via HOMER, simply run `list_homer_packages()`. For this analysis, either ‘hg38’ or ‘mm10’ packages can be installed, which include the human and mouse genomes and annotations, respectively.

~~~
## Install 'marge':
## devtools::install_bitbucket('robert_amezquita/marge', ref = 'master')
library(marge)

check_homer()

## HOMER installation found
list_homer_packages()

## # A tibble: 3x4
## status package version description
## <chr> <chr> <chr> <chr>
## 1 + homer v4.9.1 Code/Executables, ontologies, motifs for HOMER
## 2 + mouse-o v5.10 Mus musculus (mouse) accession and ontology info~
## 3 + mm10 v5.10 mouse genome and annotation for UCSC mm10
~~~

### Input data

To begin, regions of interest should first be identified. For example, peak calls derived from ATAC-seq or ChIP-seq data are of particular interest for motif enrichment analyses. At minimum, such regions should contain three columns: `chrom` (chromosome), `start`, and `end`, describing the location of the region of interest. Additional columns are also allowed - suggested columns include a region identifier, gene annotations, distance to the gene, and properties such as expression/chromatin state.

A minimal set of regions is included with the `marge` package for testing, and can be loaded as follows:

~~~
## Use included test regions from 'marge' package
test_file <- system.file('extdata/test_regions.bed', package = 'marge')
dat <- readr::read_tsv(test_file)

dat

## # A tibble: 4x6
## chrom start end id value gene
## <chr> <int> <int> <chr> <int> <chr>
## 1 chr6 102207441 102207830 region_1 1 A
## 2 chr7 44688352 44688664 region_2 2 B
## 3 chr7 92093830 92094134 region_3 3 C
## 4 chr12 103669461 103669829 region_4 4 D
~~~

### Find Motifs Across the Genome

From the regions of interest, enriched *de novo* and/or known motifs can be performed via the `find_motifs_genome()` function. This function connects to the HOMER utility `findMotifsGenome.pl`, and uses much of the same syntax, and so should be familiar to users of HOMER. The following code shows all available options, with documentation available online and via R for further details. Suffice it to say, a *de novo* and known motifs enrichment analysis is run on the test regions loaded prior, and writes results to the designated path directory (here, `results_dir`). Thus, `find_motifs_genome()` is run purely for its side-effect of creating a HOMER results directory.

~~~
## Create a temporary directory to write results
## This directory is erased once R session is closed
results_dir <- tempfile(pattern = 'test-dir_')

## Run a motif enrichment analysis
find_motifs_genome(
   dat,
   path = results_dir,
   genome = 'mm10',
   motif_length = 8,
   scan_size = 50,
   optimize_count = 2,
   background = 'automatic',
   local_background = FALSE,
   only_known = FALSE, only_denovo = FALSE,
   fdr_num = 5, cores = 1, cache = 100,
   overwrite = TRUE, keep_minimal = FALSE
)
~~~

Note in particular the argument `keep_minimal`, which if set to `TRUE`, removes all extraneous HTML and logo images, retaining only the relevant enrichment results necessary for downstream analyses. If `FALSE` (the default), the output is exactly the same as if HOMER had been run from the command line with the given parameters.

### Load Denovo and Known Motif Enrichment Results

HOMER produces two distinct files for *de novo* and known motif enrichments within the designated results directory, contained in homerMotifs.all.motif and knownResults.txt, respectively. These results can be read in using the respective `read_*_results()` functions, where `*` is either denovo or known.

~~~
known <- read_known_results(path = results_dir, homer_dir = TRUE)
<denovo <- read_denovo_results(path = results_dir, homer_dir = TRUE)

known
denovo

## # A tibble: 365 x 14
## motif~ motif~ experi~ acces~ data~ conse~ log_~ fdr tgt_~ tgt_~ bgd_~
## <chr> <chr> <chr> <chr> <chr> <chr> <dbl> <dbl> <dbl> <dbl> <dbl>
## 1 Fosl2 bZIP 3T3L1-~ GSE56~ Homer NATGA~ 1.00 1.00 1.00 0.250 2386
## 2 MafB bZIP BMM-Ma~ GSE75~ Homer WNTGC~ 1.00 1.00 1.00 0.250 2820
## 3 Fra2 bZIP Striat~ GSE43~ Homer GGATG~ 1.00 1.00 1.00 0.250 3308
## 4 Fra1 bZIP BT549-~ GSE46~ Homer NNATG~ 1.00 1.00 1.00 0.250 3747
## 5 JunB bZIP Dendri~ GSE36~ Homer RATGA~ 1.00 1.00 1.00 0.250 3900
## 6 Atf3 bZIP GBM-AT~ GSE33~ Homer DATGA~ 1.00 1.00 1.00 0.250 4570
## 7 BATF bZIP Th17-B~ GSE39~ Homer DATGA~ 1.00 1.00 1.00 0.250 4634
## 8 AP-1 bZIP ThioMa~ GSE21~ Homer VTGAC~ 1.00 1.00 1.00 0.250 5299
## 9 RUNX Runt HPC7-R~ GSE22~ Homer SAAAC~ 1.00 1.00 1.00 0.250 5731
## 10 NFY CCAAT Promot~ <NA> Homer RGCCA~ 1.00 1.00 1.00 0.250 7831
## # … with 355 more rows, and 3 more variables: bgd_pct <dbl>, motif_pwm
## # <list>, log_odds_detection <dbl>

## # A tibble: 2 x 17
## conse~ motif~ log_od~ moti~ log_p~ tgt_~ tgt_~ bgd_~ bgd_pct log_~ fdr
## <chr> <chr> <dbl> <lis> <dbl> <dbl> <dbl> <dbl> <dbl> <dbl> <dbl>
## 1 TCGCA~ 1-TCG~ 8.24 <tib~ -7.02 1.00 0.250 83.7 2.00e-4 3.00 1.00
## 2 GATTA~ 2-GAT~ 8.24 <tib~ -6.73 1.00 0.250 110 3.00e-4 2.00 1.00
## # … with 6 more variables: tgt_pos <dbl>, tgt_std <dbl>, bgd_pos <dbl>,
## # bgd_std <dbl>, strand_bias <dbl>, multiplicity <dbl>
~~~

These functions specifically are responsible for the core routine which tidies the HOMER results, and can be pointed to any existing HOMER motif enrichment result files by setting `homer_dir` to `FALSE`. The output provides a tidied version of the default HOMER output, capturing the various characteristics of enrichments, such as the identity of the motifs, target and background number/percents of the motifs across the provided sequences, statistics of enrichment, and finally the associated motif position weight matrices (PWM) in the list-column `motif_pwm`. The columns are described further in the associated help files.

### Accessing Motif PWMs

One core task of interest is working with the identified *de novo* motif PWMs that were enriched across the regions of interest. To access a specific motif PWM, the following shows several ways of accessing them from the `motif_pwm` list-column in the `known` and denovo objects previously created.

~~~
## Known Motifs
known$motif_pwm[1] *# positional access*
known$motif_pwm['OCT:OCT'] *# named access*

## Denovo Motifs
denovo$motif_pwm[1] *# positional access*
~~~

~~~
denovo$motif_pwm['1-TCGCATTG'] *# named access (denovo)*

## $'1-TCGCATTG'
## # A tibble: 8x4
## A C G T
## <dbl> <dbl> <dbl> <dbl>
## 1 0.100 0.100 0.100 0.700
## 2 0.100 0.700 0.100 0.100
## 3 0.100 0.100 0.700 0.100
## 4 0.100 0.700 0.100 0.100
## 5 0.700 0.100 0.100 0.100
## 6 0.100 0.100 0.100 0.700
## 7 0.100 0.100 0.100 0.700
## 8 0.100 0.100 0.700 0.100
~~~

Note the column names identifying the nucleotide, the rows for the ordered nucleotide position, and the frequencies at each cell, where each row sums to 1.

### Writing Motif PWMs - Manual

To write a motif to a file, the helper function `write_homer_motif()` can be used, and values can be manually specified for motifs of interest.

~~~
## Note double bracket to access list value;
## single bracket produces error
write_homer_motif(
    motif_pwm = denovo$motif_pwm[['1-TCGCATTG']],
    motif_name = 'my_first_motif',
    log_odds_detection = 4.35,
    consensus = 'CACATCCT',
    file = paste0(results_dir, '/my_first_motif.motif')
)
~~~

### Writing Motif PWMs - Automatic

Oftentimes, one wants to write all associated motif PWMs from a given *de novo* enrichment search. Given that `write_homer_motif` produces a single motif at a time, we need a way to produce a file for each row of the denovo object.

While there are various ways to iterate over the rows of the denovo object, such as with `for-loops` or the `apply` family of functions, below is shown a method using the `purrr` package from the `tidyverse` family, using the `map()` and `walk()` functions. Briefly, these functions apply a function to each element of a vector, thus allowing for iteration over the rows of an object such as those we created, `known` and denovo. `map` returns the transformed object, whereas `walk` calls the function for its side-effect, such as in the case of writing a file as in `write_homer_motif()`. Given that we need several varying inputs (all the arguments of `write_homer_motif()`, we use the variant `pwalk()` to allow for our many varying inputs. The resulting code below writes all of our denovo results (three motifs in total) to unique files in our temporary `R` results directory.

~~~
## Write to multiple files - vary the ' file' argument
library(purrr)
pwalk(
      .l = list(
         motif_pwm = denovo$motif_pwm,
         motif_name = denovo$motif_name,
         log_odds_detection = denovo$log_odds_detection,
         consensus = denovo$consensus,
         file = paste0(results_dir, '/', denovo$motif_name, '.motif')
     ),
      .f = write_homer_motif
)

## Write to a single file with all motifs
## Keep the 'file' argument constant
my_motifs_file <- paste0(results_dir, '/my-motifs.motif')

## Append by default is TRUE, e.g. can write more than one
## motif at once, but this means don't run this code more
## than once!
pwalk(
     .l = list(
        motif_pwm = denovo$motif_pwm,
        motif_name = denovo$motif_name,
        log_odds_detection = denovo$log_odds_detection,
        consensus = denovo$consensus
     ),
     .f = write_homer_motif,
     file = my_motifs_file,
     append = TRUE
)
~~~

### Search for Instances of Motifs Across Regions

Now that we have produced our motif PWM as a file, it can be used to map the motif back to where it occurs in our set of regions using the `find_motifs_instances()` function. While it is also a wrapper for `findMotifsGenome.pl`, it partitions out a unique piece of functionality that presents an entirely different type of results. Note that this function *requires* the use of a HOMER motif file that is written out (either by hand or by the `write_homer_motif()` function) as it contains the necessary parametrization for motif finding.

Also note that we can search for either a single motif or multiple motifs concurrently, where the multiple motifs are written into a single file, as shown above. Here, we will search for all three of the *de novo* motifs identified initially in our regions.

~~~
motif_instances_file <- paste0(results_dir, '/motif-instances.txt')

find_motifs_instances(
    dat,
    path = motif_instances_file,
    genome = 'mm10',
    motif_file = my_motifs_file,
    scan_size = 50,
    cores = 1, cache = 100
)
~~~

This function, similarly to `find_motifs_genome()` and `write_homer_motif()`, produces output that is written to into a file outside of `R`, and similarly relies on a helper function to read the resulting output.

### Read Instances of Motifs Across Regions

The function `read_motifs_instances()` works in tandem with `find_motifs_instances()` to read its output, returning the location and scores of the specified motifs from the previous step.

~~~
motif_instances <- read_motifs_instances(motif_instances_file)

motif_instances

## # A tibble: 2x6
## region_id offset sequence motif_name strand motif_score
## <chr> <int> <chr> <chr> <chr> <dbl>
## 1 region_2 -24 TCGCATTG 1-TCGCATTG + 8.24
## 2 region_1 -5 GATTACTC 2-GATTACTC + 8.24
~~~

### Advanced Usage

Using the verbs learned in the previous section, this part will focus on tying it all together to create a framework for testing for motif enrichment and subsequently analysing the results across multiple sets of regions in a tidy fashion. This section relies heavily on the tidyverse framework, and in particular will use the tools from `purrr, map()/walk()`, introduced in the previous section. Additionally, we will simulate a dataset using the `valr` package function `bed_random()`. While outside of the scope of this tutorial, valr provides additional functionality to work with genomic intervals using similar `tidyverse` conventions, and is recommended for manipulating regions of interest from high-throughput experiments.

Below are the libraries required for this next section.

~~~
## Required libraries for advanced usage tutorial
library(valr)
library(dplyr)
library(tidyr)
library(purrr)
library(ggplot2)
~~~

### Constructing Multiple Simulated Sets of Regions

By bringing the usage of HOMER into `R` via an API, it becomes possible to construct multiple queries, execute them all, and then read in the results from all the queries.

First, we will create a simulated set of intervals for which to test for motif enrichment, formatting the regions into a `tibble` object which allows for the creation of list-columns, which tidily encapsulate the genomic intervals.

~~~
## 1. Construct multiple region samples ‐‐‐‐‐‐‐‐‐‐‐‐‐‐‐‐‐‐‐‐‐‐‐‐‐‐‐‐‐‐‐‐‐‐‐‐‐
## Create genome from first two chromosomes of mouse
genome <- data.frame(chrom = c('chr1', 'chr2'),
                            size = c(195471971, 182113224))

## Set seeds and ids for the simulated sets of regions
seed <- 1:5
id <- letters[1:5]

## Simulate different sets of regions (each with unique seeds) and munge
## into the tibble format, keeping the regions as a list-column
sim <- map(seed, ~ valr::bed_random(genome, length = 50, n = 50, seed = .x))
tbl_regions <- tibble(id = id, regions = sim)

## Inspect overall tibble organization
tbl_regions

## # A tibble: 5x2
## id regions
## <chr> <list>
## 1 a <tibble [50 x 3]>
## 2 b <tibble [50 x 3]>
## 3 c <tibble [50 x 3]>
## 4 d <tibble [50 x 3]>
## 5 e <tibble [50 x 3]>
## View regions sample
tbl_regions$regions[[1]] %>% print(n = 5)

## # A tibble: 50 x 3
## chrom start end
## <chr> <int> <int>
## 1 chr1 3322213 3322263
## 2 chr1 12952049 12952099
## 3 chr1 13419401 13419451
## 4 chr1 13496953 13497003
## 5 chr1 44639848 44639898
## # … with 45 more rows
~~~

### Finding Motifs Across Multiple Sets of Regions

The next step involves specifying the paths to the results directory for each respective set of regions. This is necessary for `find_motifs_genome()` to output the motif enrichment results for each set of regions separately (otherwise the results will be overwritten multiple times).

~~~
## 2. Find Motifs in Genome ‐‐‐‐‐‐‐‐‐‐‐‐‐‐‐‐‐‐‐‐‐‐‐‐‐‐‐‐‐‐‐‐‐‐‐‐‐‐‐‐‐‐‐‐‐‐‐‐‐

## Create new (temp) results directory
results_dir <- tempfile(pattern = 'ht-dir_')

## Append results path to tibble of regions
tbl_regions <- tbl_regions %>%
    mutate(path = paste0(results_dir, '/homer_', id))
tbl_regions[1, ] *# inspect resulting first row*

## # A tibble: 1x3
## id regions path
## <chr> <list> <chr>
## 1 a <tibble [50 x 3]> /var/folders/nj/s_mzzyn922l8vzmll72t0vdr0000gn/~
~~~

Now, using each set of regions and the unique paths for each set of regions, `find_motifs_genome()` can be run, varying across these two parameters, and setting as constant the various motif enrichment search parameters for all the different sets.

One other potential application not shown here is testing how these various parameters affect motif enrichment results if we were instead to keep the regions tested as constant instead. This is left as an exercise for the interested reader.

~~~
## Iterate over all regions to perform motif finding
## Writing each set of regions results to different directory
## Only perform known motif results enrichments
pwalk(
     ## Varying parameters (regions, output directory)
     .l = list(x = tbl_regions$regions, path = tbl_regions$path),
     ## Function to find motifs across genome
     .f = find_motifs_genome,
     ## Constant parameters (motif search settings)
     genome = 'mm10',
     scan_size = 50,
     optimize_count = 2,
     only_known = TRUE,
     cores = 2, cache = 100,
     keep_minimal = TRUE, overwrite = TRUE
)
~~~

### Combining Results from Multiple Motif Enrichment Searches

Following the motif enrichment search for each set of regions, the results can be read in, iterating over the variable `path` within the `tbl_regions` object to pull in the relevant results. The results are cleaned to remove the `path` and `regions` columns subsequently, and then expanded such that each individual motif enrichment result has its corresponding id as a separate variable.

~~~
## 3. Read in All Results ‐‐‐‐‐‐‐‐‐‐‐‐‐‐‐‐‐‐‐‐‐‐‐‐‐‐‐‐‐‐‐‐‐‐‐‐‐‐‐‐‐‐‐‐‐‐‐‐‐‐‐‐

## Read known motif enrichment results into a new list-column
## Remove the path and regions columns for cleanliness
## Unnest results to expand into plotting friendly format
tbl_results <- tbl_regions %>%
     mutate(known_results = map(path, read_known_results)) %>%
     select(-path,-regions) %>%
     unnest()
~~~

~~~
## View sampling of results
tbl_results %>% sample_n(size = 5)

## # A tibble: 5 x 15
## id motif~ motif~ experim~ acce~ data~ conse~ log_~ fdr tgt_~ tgt_p~
## <chr> <chr> <chr> <chr> <chr> <chr> <chr> <dbl> <dbl> <dbl> <dbl>
## 1 e Nr5a2 NR Pancrea~ GSE3~ Homer BTCAA~ 1.00 1.00 3.00 0.0638
## 2 d Pit1 Homeo~ GCrat-P~ GSE5~ Homer ATGMA~ 0 1.00 2.00 0.0408
## 3 b E2F4 E2F K562-E2~ GSE3~ Homer GGCGG~ 0 1.00 0 0
## 4 a NPAS bHLH Liver-N~ GSE3~ Homer NVCAC~ 0 1.00 4.00 0.08001
## 5 b ZNF317 Zf HEK293-~ GSE5~ Homer GTCWG~ 0 1.00 0 0
## # … with 4 more variables: bgd_num <dbl>, bgd_pct <dbl>, motif_pwm
## # <list>, log_odds_detection <dbl>
~~~

### Summarising Characteristics of Motif Enrichment Results

In this tidy form, it becomes possible to summarise results on the basis of motif families, as opposed to simply individual motifs. Below, we use the `id` and `motif_family` columns to group the results, and summarise a given motif family by taking the most significant individual motif appearance as the summary statistic (e.g., the max-log10 p-value).

~~~
## Summarise number of
tbl_summary <- tbl_results %>%
     group_by(id, motif_family) %>%
     summarise(top_log_p_value = max(log_p_value)) %>%
     ungroup()

## Print example results
tbl_summary %>% sample_n(5)

## # A tibble: 5x3
## id motif_family top_log_p_value
## <chr> <chr> <dbl>
## 1 c CCAAT 0
## 2 c Zf? 0
## 3 a ? 0
## 4 c EBF 0
## 5 d Forkhead 0
~~~

Note that few motifs appear significant in this simulated dataset, as regions were chosen at random, without a true biological correlate.

### Plotting Results

Finally, we can inspect the results of our analyses using `ggplot2` package conventions, given the tidiness of the results. Here we show a subset of the overall results for succinctness.

~~~
families <- c('AP2', 'bHLH', 'bZIP', 'CCAAT', 'CP2')

tbl_summary %>%
     filter(motif_family %in% families) %>%
     ggplot(aes(x = motif_family, y = id, fill = top_log_p_value)) +
     geom_tile()
~~~

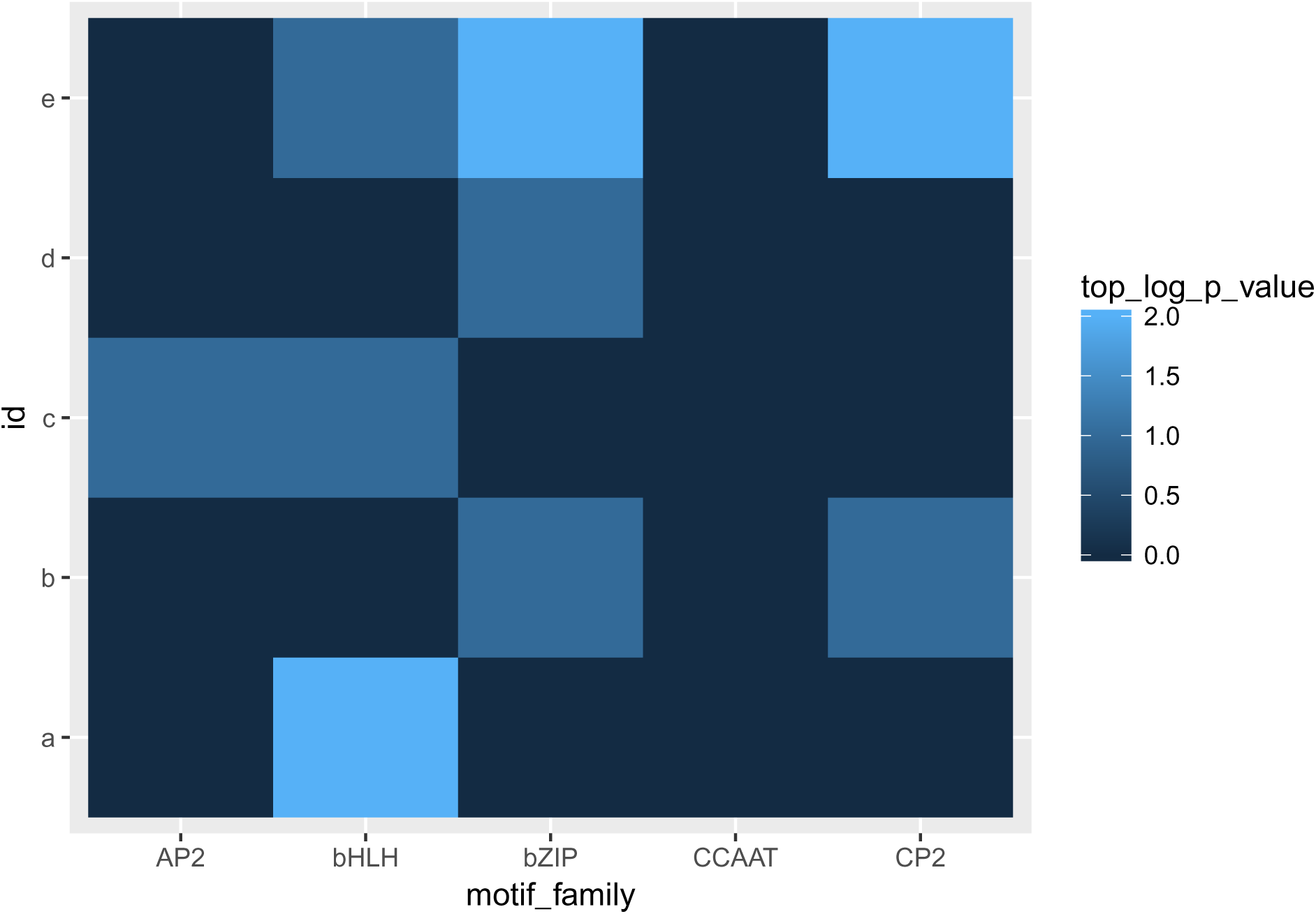

## Summary

`marge` presents an `R` centric form of performing and analyzing motif enrichment results using the popular HOMER suite of tools, providing an easy-to-use interface and results in line with modern `R` idioms. We envision that `marge` will serve as a valuable tool to assist researchers in performing motif enrichment analyses quickly, easily, and reproducibly.

## Data and Software Availability

Online documentation for HOMER and installation details can be found at: http://homer.ucsd.edu/homer/ `marge` can be installed via devtools using: `devtools::install_bitbucket('robert_amezquita/marge', ref = 'master'`)

The latest `marge` source code is available at: https://bitbucket.org/robert_amezquita/marge/

Stable versions of marge are located in the Downloads section of the source code repository, under the tab ‘Tags’, at: https://bitbucket.org/robert_amezquita/marge/downloads/

Online documentation for `marge` can be found at: https://marge.aerobatic.io/

## Acknowledgements

R.A.A. wrote the vignette, designed and implemented the software package. Jason Vander Heiden edited the manuscript and reviewed source code for clarity. Greg Finak contributed to code sussing out HOMER installation whereabouts.

